# Pain sensitivity and athletic performance

**DOI:** 10.1101/514224

**Authors:** Lior Zeller, Nadav Shimoni, Alina Vodonos, Iftach Sagy, Leonid Barski, Dan Buskila

**Affiliations:** Department of Medicine C, Soroka University Medical Center; Faculty of Health Sciences, Beer-Sheva, Israel; Ben-Gurion University of the Negev, Beer-Sheva, Israel; Clinical Research Center, Soroka University Medical Center; Faculty of Health Sciences, Beer-Sheva, Israel; Department of Medicine F, Soroka University Medical Center; Faculty of Health Sciences, Beer-Sheva, Israel; Department of Medicine H, Soroka University Medical Center; Faculty of Health Sciences, Beer-Sheva, Israel

**Keywords:** Pain threshold, Long distance runners, SF-36, Cold pressor test, athletic performance, pain tolerance

## Abstract

**Purpese:** To determine whether higher pain thresholds are associated with better performance in long-distance runners.

**Design:** Cross-sectional study.

**Methods:** Seventy participants, divided into groups of fast and non-fast runners according to peak results in a 10km run. Main Outcome Measures, Cold pressor test.

**Results:** Of the 70 subjects, 28 were in the fastest group (less than 39 minutes in a 10km run) and 42 in the non-fast group. The faster group was characterized with older age (34.0±8.5 vs. 29.5±5.7, *p*=0.01), greater mean weekly running time (5.5 (0-17) vs. 2 (0- 10), *p*<0.001), and more years of running [10 (1.5-34.0) vs. 7 (0-20, *p*=0.05)]. In a multivariable analysis longer cold pressor time was associated with faster 10Km run (O.R 1.01, 95% C.I 1.00-1.01).

**Conclusions:** It seems that higher pain thresholds play an important role in the superior ability of long distance runners.

## Introduction

After setting the world record for women in a marathon run (2:15:25) at the London Marathon in 2003, a record that has not yet been broken, Paula Radcliffe was asked what helped her to achieve such an impressive result. Her answer was that it was in her ability to tolerate pain better and thereby break her own “boundaries” again and again^1^.

Pain perception is investigated using a variety of tests and questionnaires, for example the cold pressor^2^ test, and the MOS SF-36 questionnaire for health-related quality of life assessment.^3^ The association between physical training and pain perception has been studied mainly by comparing trained and untrained populations. On the one hand, it appears that professional physical training in endurance sports causes a change in pain perception^4^ On the other hand, high pain threshold and high pain resistance may lead certain people to take part in this kind of physical activity.^5^

There is evidence of a link between incidence of pain, the impact of pain on quality of life, and even the efficacy of treatment of pain-related situations and belonging to a particular ethnic group. For example, it has been shown that an individual belonging to an ethnic minority in his country is likely to have higher pain tolerance than other populations in that country. Hence, it appears that there is a difference in pain perception between different populations.^6^

However, the link between runners’ performance and their pain tolerance threshold or their ability to tolerate pain remained unclear. We hypothesize that higher pain thresholds and tolerance may lead to better performance among long-distance runners and can serve as a predictor of running ability among runners. Consequently, we expect that faster runners will demonstrate greater tolerance in the cold pressor test.

## Methods

We conducted a cross-sectional study of fast and non-fast runners. The participants were divided in accordance with their best result in a 10km run. Runners with a result of 39 minutes or less on a 10km run were assigned to the fast group, while those who achieved a result over 39 minutes but less than 50 minutes were assigned to the non-fast group. Division into groups according to a 39-minute threshold was based on an analysis of the distribution of the results of 10km races in Israel, which revealed that a result faster than 39 minutes was included in the top quintile. Inclusion criteria for the study were male gender, age over 18 years, membership in a running club or an individual training program, absence of injuries at the time of initiation of the study, and absence of chronic diseases. Exclusion criteria included subjects with Raynaud’s syndrome, as they aren't eligible for the cold pressor test. This study was approved by the institutional ethics committee.

Each participant was asked to fill out a socio-demographic questionnaire and note his running results for different distances. In order to assess pain perception and threshold, we used the SF36 MOS questionnaire and the cold pressor test.

The cold pressor test is based on neutral stimulation that activates pain receptors to cold and is used to test for pain tolerance^2^. Sensitivity to cold-induced pain has been proven to be related to sensitivity to mechanical pain, and high tolerance for the cold pressor may be associated with inhibition of pain activation from a central source. Several studies have demonstrated the diagnostic ability of this test, for example documenting relatively high pain tolerance among professional dancers and relatively low tolerance among patients with chronic low back pain^7^.

In the framework test, the subjects placed one of their arms up to the elbow into a bucket with water at a constant temperature of about 4°C and reported pain level every 10 seconds on a scale of 1 to 10 (max.). The test lasted 4 minutes, or alternatively was discontinued if the subject reported maximum pain level (10). The length of time the participants left their arms in the water was measured in seconds by a stopwatch and was recorded^8,9^.

Data are expressed as mean ± standard deviation (SD) and median ± inter-quartile range (IQR). We compared runners' characteristics using t-test and Mann-Whitney tests. We conducted a multivariable logistic regression model to examine whether higher cold pressor test results are associated with belonging to the 10Km fast group. The final model was selected according to the statistical significance of coefficients, their clinical relevance, and the model discriminatory characteristic, which were evaluated by calculating the c-statistic (statistics showing the discriminatory power of the model, i.e., difference in the predicted probabilities of the event between those with and without the event) and by Hosmer Lemeshow in addition to choosing the minimal −2 log likelihood of each model (which represent the goodness of fit for binary outcomes in a logistic regression). In all the tests, a significance level of *p* < 0.05 was determined. Data analysis was performed with SPSS version 21.

## Results

Seventy subjects were found to fit the inclusion criteria (men over age 18 with a 10km run result under 50 minutes) and agreed to take part in the study. Twenty-eight (28) subjects were assigned to the fast group (10km in 39 minutes or less) and 42 to the nonfast group. All of the subjects took part in the entire study in accordance with the study regulations and filled in the socio-demographic questionnaire, the SF-36 Questionnaire for Quality of Life Evaluation and underwent the cold pressor test.

Table 1 presents the characteristics of the two study groups. There were a number of statistically significant differences between the fast and the non-fast groups: older age (34.0±8.5 vs 29.5±5.7, *p* = 0.01), greater median weekly running time [5.5 (0-17) vs. 2 (0-10), *p* < 0.001], and more years of running significance [10 (1.5-34.0) vs. 7 (0-20), *p* = 0.05]. In the analysis of the SF-36 questionnaire, no statistically significant difference was found between the two groups for all components of the questionnaire.

**Table 1.**
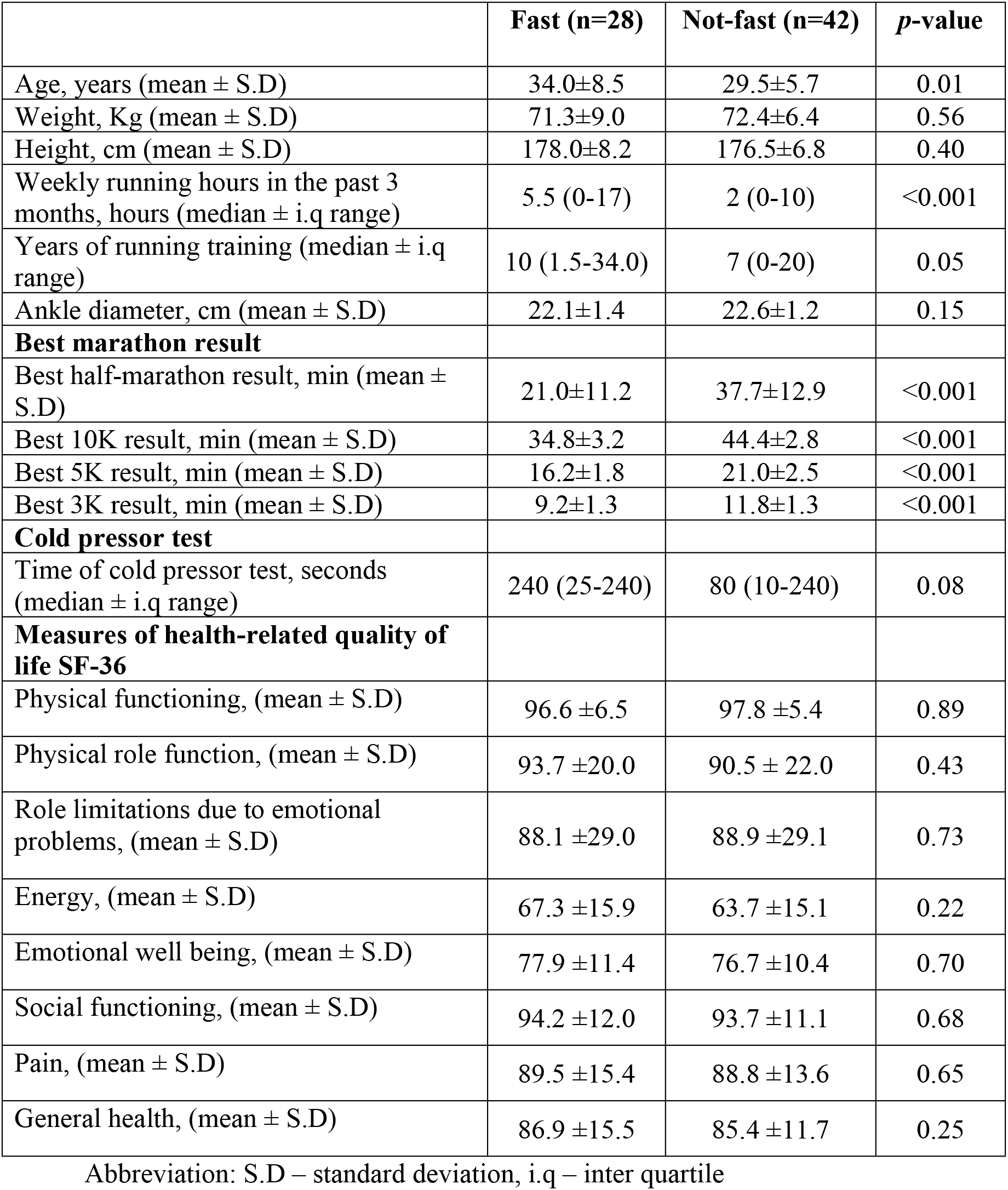
Subjects’ characteristics.

We found a marginal difference of test time between the two groups, as shown in Table 1 (240 seconds in the fast group vs. 80 seconds in the control group, *p* = 0.08). It should be noted that 240 seconds is the maximum test time, i.e., the median time in the fast group is also the maximum time of the test. Further we performed a multivariate logistic regression model while controlling for additional possible confounders associated with belonging to the faster 10Km group, as shown in Table 2. After adjusting for running experience (in years) and the age of the participant we found that higher cold pressor results remained significant in association with being in the fast group (O.R 1.01. 95% C.I 1.00-1.01).

**Table 2.** Logistic regression for predicting inclusion in the fast running group.

| Variable | OR | 95% CI | *p*-value |
| --- | --- | --- | --- |
| Cold pressor total time | 1.01 | 1.00-1.01 | 0.04 |
| Years of running | 1.01 | 1.01-1.19 | 0.04 |
| Age | 1.07 | 0.98-1.17 | 0.11 |

## Discussion

In this study we examined the association between pain tolerance and the performance of long-distance runners based on the assumption that a higher pain threshold and greater pain tolerance for longer periods may contribute to better running performance in these runners. To the best of our knowledge, this assumption has not yet been examined previously. Our main finding is that pain threshold and tolerance is associated with better performance of long distance runners. Our results remained consistent even in multivariable analysis.

The possible reasons for the superior ability of specific groups of runners (e.g. Kenyan and Ethiopian runners) has created a lively focus for discussion and a number of hypotheses attempting to explain this phenomenon have emerged over the years, among them superior genetics, high hemoglobin levels and hematocrit, metabolism and muscle economy enhanced by a unique lower limb structure (ankle circumference and muscle structure), living conditions, training at high altitude, and so on^10,11,12^. However, despite these attempts, no specific factor has been proven to explain the superior performance of the East African runners.

According to the literature, there are two significant factors that determine the performance of a long-distance runner:

1. The lactic acid threshold (the “anaerobic threshold”, the level of activity of the muscle in which the lactic acid production rate exceeds the evacuation rate) is a trait that can be improved by training^13^.
2. The maximum oxygen consumption (VO_2_MAX) and the athlete’s performance at this point, a feature almost entirely determined at birth, was also proposed as a variable explaining the control of East Africans. ^14,15,16^

For very good reasons, the field of long-distance running evokes in those who follow it the sense that there are factors beyond time and quality of training that help certain runners excel in comparison to others. A small group of East African runners dominates these races almost unchallenged. At the same time, and despite the fact that a considerable number of factors that might explain this phenomenon have been investigated, until now, none of them has been proven to be significantly related to better running performance other than, of course, quality training^11,12^.

The relationship between pain perception and sports has been studied mainly as a factor that distinguishes between a population that engages in sports and one that does not for the purpose of examining the effect of exercise on pain perception^11^ or on pain threshold and endurance as predictors of the choice to engage in physical exercise.^5,7^ In addition, it has been shown that there is a difference in pain perception between different populations (e.g children and adolescents)^2^.

This cross-sectional study compared a group of fast runners and a group of nonfast runners, divided according to their top scores in a 10km run. An analysis of the results demonstrated that there were a number of statistically significant differences between the fast group and the non-fast group. The fast runners were older, with more experienced with running, and they practiced more during the week. However, there was no significant difference in ankle circumference between the groups, despite the fact that this variable has been suggested in the literature as a predictor of better long-distance running ability^11^.

In order to test pain threshold and tolerance, we chose to use the cold pressor test and the SF 36 test, taking into account that pain is not an objective measure and there are different types of pain. The cold pressor test is used in the literature for comparison between athletes and non-athletes^2,7^. In contrast the SF 36 test is used mainly to compare healthy and non-healthy populations^3^.

In examining the results of the cold pressor test, several interesting points became apparent. First, the attempt to test the existence of a linear relationship between the cold pressor results (i.e., higher pain threshold) and running results demonstrated that no such relationship exists. This is true for both the fast and the non-fast groups. However, it was found that the group of fast runners had a higher score than the non-fast runners at a marginal level of statistical significance. In addition, a longer period of time in the cold pressor test has been shown to predict belonging to the fast group, with each additional test unit increasing the chance times. Given this, and based on the assumption that a longer period of time in the cold pressor test reflects higher pain threshold and tolerance^2,7^, an association between higher pain threshold and greater tolerance to pain and running ability may be inferred (as it is expressed in a 10km run).

The analysis of the SF-36 questionnaire did not show statistically significant differences between groups. This may be due to the fact that this questionnaire is designed to evaluate patients with some kind of illness, in contrast to this study’s subjects, who were all healthy and physically active (including those in the non-fast group).

We acknowledge several limitations of this study. It seems that many parameters can affect the results of the cold pressor test as it represents a subjective measurement of pain, especially given the competitive nature of all of the subjects. As a result, at the time of the tests, we eliminated several parameters that could create a bias in the results such as preventing the presence of an additional person with the subject at the time of the test, and performed the tests in similar weather conditions, using a similar predefined set of orders given to the subjects. Although we placed a great deal of emphasis on creating uniformity among the subjects in the performance of these tests, it is possible that there are additional parameters that were not taken into account and may have created a bias in the test results. In addition, most of the study population consisted of Israeli-born runners. It is possible that in an analysis of other nationalities, particularly East Africans, known for their excellence in long runs, other data would have emerged. Finally, most of the runners in the fast group were characterized by older age, many years of training, and significantly more weekly running hours (weekly training time) than the non-fast group. However, the multivariable analysis was adjusted for these variables in order to minimize their possible confounder effect.

## Conclusion

Our findings imply that higher pain threshold and tolerance may be associated with better achievements of long distance runners. To the best of our knowledge, this study is the first to clearly delineate this relation. Although previous studies which focused on other physiological variables (e.g. anaerobic threshold and VO_2_MAX) showed poor association with improved performances, our study provide opportunities for future research in the field of pain perception among elite athletes.

## Abbreviations

CI: Confidence interval
IQR: Interquartile range
OR: Odds ratio
SP: Standard deviation

